# Laboratory evolution of synthetic electron transport system variants reveals a larger metabolic respiratory system and its plasticity

**DOI:** 10.1101/2022.04.04.487013

**Authors:** Amitesh Anand, Arjun Patel, Ke Chen, Connor A. Olson, Patrick V. Phaneuf, Cameron Lamoureux, Ying Hefner, Richard Szubin, Adam M. Feist, Bernhard O. Palsson

## Abstract

Respiration requires organisms to have an electron transport system (ETS) for the generation of proton motive force across the membrane that drives ATP synthase. Although the molecular details of the ETS are well studied and constitute textbook material, few studies have appeared to elucidate its systems biology. The most thermodynamically efficient ETS consists of two enzymes, an NADH: quinone oxidoreductase (NqRED) and a dioxygen reductase (O_2_RED), which facilitate the shuttling of electrons from NADH to oxygen. However, evolution has produced variations within ETS which modulate the overall energy efficiency of the system even within the same organism ^1–3^. The system-level impact of these variations and their individual physiological optimality remain poorly determined. To mimic varying ETS efficiency we generated four *Escherichia coli* deletion strains (named ETS-1H, 2H, 3H, and 4H) harboring unbranched ETS variants that pump 1, 2, 3, or 4 proton(s) per electron respectively. We then used a combination of synergistic methods (laboratory evolution, multi-omic analyses, and computation of proteome allocation) to characterize these ETS variants. We found that: (a) all four ETS variants evolved to a similar optimized growth rate, (b) the evolution of ETS variants was enabled by specific rewiring of major energy-generating pathways that couple to the ETS to optimize their ATP production capability, (c) proteome allocation per ATP generated was the same for all the variants, (d) the aero-type, that designates the overall ATP generation strategy ^4^ of a variant, remained conserved during its laboratory evolution, with the exception of the ETS-4H variant, and (e) integrated computational analysis of then data supported a proton-to-ATP ratio of 10 protons per 3 ATP for ATP synthase for all four ETS variants. We thus have defined the Aero-Type System (ATS) as a generalization of the aerobic bioenergetics, which is descriptive of the metabolic systems biology of respiration and demonstrates its plasticity.

---

*E. coli* has a highly flexible ETS consisting of 15 dehydrogenases and 10 reductases to allow growth in both oxic and anoxic environments ^5^. The expression of these enzymes is regulated by a variety of electron acceptors with a known hierarchy, such that oxygen represses all anoxic respiratory pathways and nitrate represses other anoxic pathways ^3,5^. Despite this thermodynamic hierarchy, co-expression of different respiratory chains was reported in another γ-proteobacterium to expand the flexibility of its electron transfer network ^3^. We probed the condition-dependent expression of all these dehydrogenases and reductases using a large RNA-seq compendium for *E. coli* ^6^. We observed a spectrum of expression values of these genes across the experimental conditions showing the contribution of these enzymes in generating plasticity in energy metabolism (Supplementary Figure 1).

To examine the contributions of individual oxic respiratory pathways to bioenergetics, we sought to design unbranched pathways through the ETS. The oxic component is contributed by both proton pumping and non-pumping NqREDs (hereafter referred to as NDH-I and NDH-II, respectively) along with three types of O_2_REDs (Figure 1A). Cytochrome bd O_2_REDs (CBDs) are less electrogenic compared to Cytochrome bo_3_ O_2_REDs (CYO). There are two CBDs, bd-I and bd-II, and both function similarly to generate proton-motive force (PMF) by a scalar movement of protons involving transmembrane charge separation. The similar PMF generation strategies make bd-I and bd-II O_2_REDs equivalent when choosing gene knock-out strategies ^7,8^.

**Figure 1:**
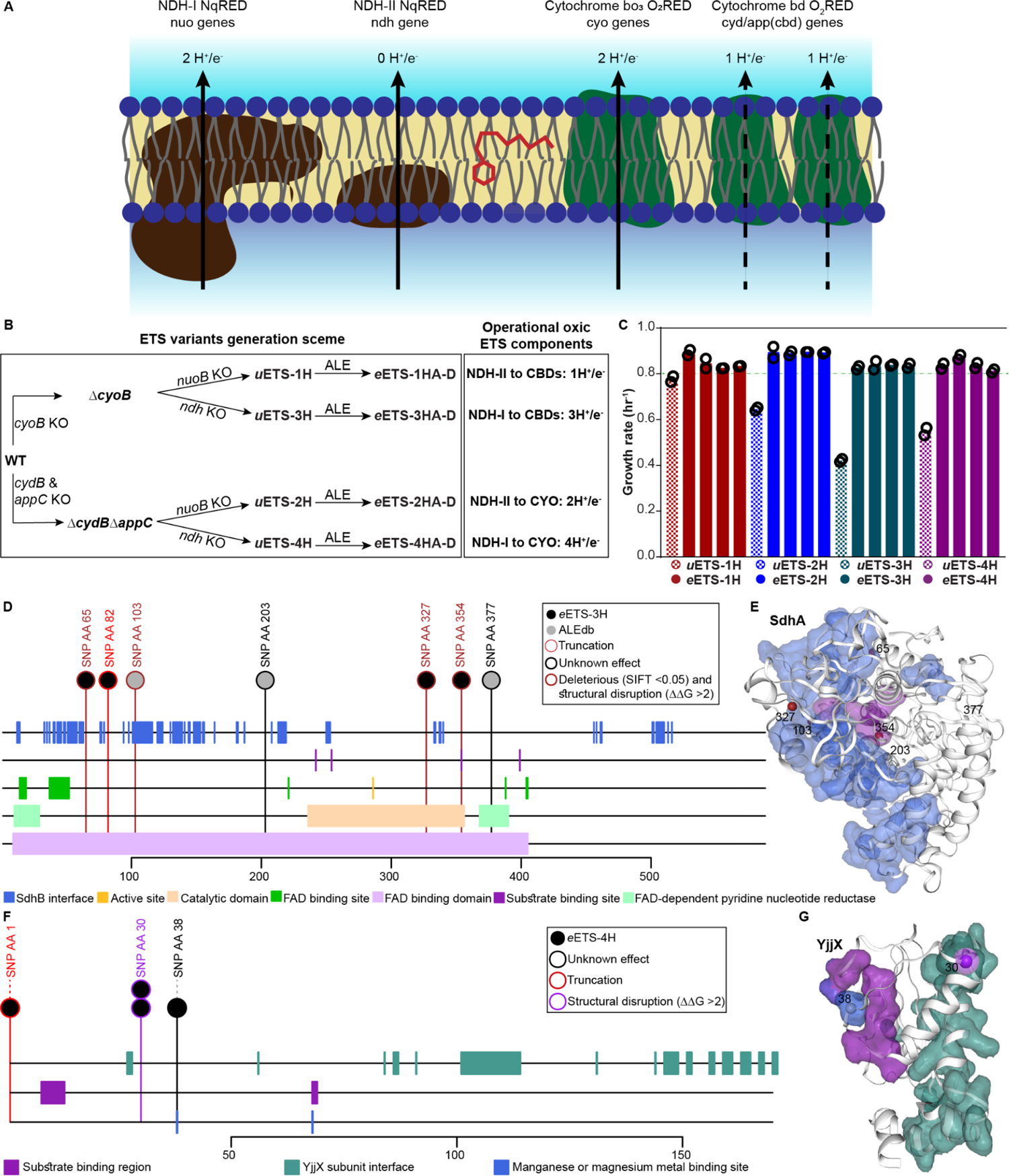
Generation and evolution of unbranched ETS variants: (A) Schematic showing the four respiratory enzymes involved in the flow of electrons from NADH (donor) to oxygen (acceptor). NDH-I and NDH-II are the proton pumping and non-pumping NADH: quinone oxidoreductase, respectively. Dashed arrows for CBDs represent the scalar mode of PMF generation. (B) Scheme for generating ETS variants translocating 1, 2, 3, or 4 proton(s) per electron. *u*ETS is the unevolved strain and *e*ETS is the evolved strain. A-D are the four independently evolved lineages of each strain. (C) Growth rates of ETS variants before and after ALE. (D, F) Predictive mechanistic interpretation of the impact of mutations observed in the evolved strains of (D) ETS-3H (*sdhA*) and (F) ETS-4H (*yjjX*) mutations on the structure and function of the protein. Mutations displayed are those from this study and other ALE experiments in ALEdb seen to mutate these genes ^9^. Horizontal tracks display the reported features associated with the region of the protein. The mutations collected from ALEdb refer to experiments from the following set, respectively ^10–12^. Protein structures showing the amino acid residues mutated in (E) SdhA and (G) YjjX.

Based on these characteristics, we designed four ETS variants with unbranched electron flows representing all alternate oxic respiratory routes translocating 1, 2, 3, or 4 proton(s) per electron (designated as ETS-nH, with n=1,2,3,4). The designs of the four ETS variants are illustrated in Figure 1B.

Next, we analyzed their growth phenotype (Figure 1C). Interestingly, the unevolved variants (called *u*ETS) showed different growth rates that had no clear association with their H^+^/e^-^ value. While the loss of activity of NDH-I showed a lesser growth rate retardation, the deletion of NDH-II significantly compromised the growth rate of the deletion strains.

To allow the ETS variants to overcome the growth defects resulting from gene deletions, we performed adaptive laboratory evolution (ALE) with four independent replicates of each variant in an oxic environment (evolved variants are named *e*ETS-nHm with the replicate evolutionary endpoints indexed as m = A, B, C, D) ^13^. We evolved all variants until their growth rate plateaued. ETS-1H, 2H & 3H required evolution for approximately 400 generations, while ETS-4H required approximately 700 generations. In spite of the different number of protons pumped per electron, all four ETS variants evolved to a similar optimized growth rate in replicate evolutions (∼0.85 hour^-1^) (Figure 1C).

Next, we sought to determine the acquired mutations that enabled adaptation to a higher growth rate for all ETS variants. We performed whole-genome sequencing of each strain and used a comprehensive database of mutations from ALE experiments (aledb.org ^9^) to interpret the potential impact of the identified mutations. The mutation calling revealed only a few genetic changes in the evolved strains (1-6) except for *e*ETS-4HC which acquired 15 genetic changes. The higher number of mutations in *e*ETS-4HC could be due to the mutated DNA mismatch repair enzyme *mutS* in this strain. An intergenic mutation between *pyrE* and *rph* has been reported to alleviate pyrimidine pseudo-auxotrophy resulting in a faster growth rate. RNA polymerase subunit mutations are proposed to favor a higher growth rate by accelerating the transcriptional processes. Another common mutation reported to support a faster growth rate is in the intergenic region between *hns* and *tdk*. This mutation is expected to downregulate several stress response pathways and shift resources to support growth. Every ETS variant acquired mutations responsible for enabling faster growth on M9 minimal medium (Supplementary Table 1 ^10,14–16^. *u*ETS-1H carried the *pyrE-rph* intergenic mutation, which explains the relatively faster initial growth rate of this strain.

Besides mutations responsible for acclimatization to media, *u*ETS-3H and *u*ETS-4H acquired a common gene-related mutation in all four independently evolved lineages. This mutational convergence simplified the otherwise difficult task of establishing the genotype-phenotype relationship ^17–19^.

All four evolved replicates of *u*ETS-3H acquired point mutations in *sdhA*, the catalytic subunit of succinate dehydrogenase (Supplementary Table 1). *e*ETS-3HB acquired a point mutation that brings in a premature termination codon in the *sdhA* open reading frame, suggesting a loss of functional enzyme. We explored the potential impact of other mutations by investigating whether the SNPs could affect the protein’s function based on amino acid properties and sequence homology (SIFT) ^20^ or structural stability (ΔΔG) ^21^. Almost all mutations in *sdhA* were either in or near interface surfaces and seem to be working to disrupt its functionality by either disrupting a substrate-binding site or causing a structural-functional perturbation (Figure 1D & E). Notably, the deletion of another subunit of this enzyme, *sdhC*, has been reported to increase the biomass yield in an oxic environment ^22^. *u*ETS-3H appeared to adopt a similar metabolic route to increase its growth rate.

All four replicates of *u*ETS-4H acquired mutations in an inadequately characterized gene, *yjjX* (Supplementary Table 1, Supplementary Table 1). The structural and biochemical evidence suggests that YjjX, an inosine/xanthosine triphosphatase, may be involved in the mitigation of the deleterious impact of oxidative stress by preventing the accumulation of altered nucleotides ^23^. Also, the physical association of YjjX with the elongation factor suggests a negative impact on the translational rate. The STRING-based protein-protein interaction predicted the association of YjjX with glycolytic and ATP biosynthetic processes ^24^. Interestingly, *e*ETS-4HA replaced the start codon, ATG, with ATA. Apart from replacing methionine with isoleucine, this substitution potentially diminishes the expression of this protein ^25^. A similar disruptive impact is expected from other *yjjX* mutations (Figure 1F & G). The cysteine to tyrosine substitution at amino acid residue 30 was predicted to destabilize the structure as it lies just beside a subunit interface residue, and a charge reversion due to the glutamate to lysine substitution at amino acid residue 38 targets the metal-binding site. Thus, it appears that *e*ETS-4H is attempting to prevent translational halting to achieve a higher growth rate.

Since the restoration of the evolved variants to the same growth rate cannot be deciphered from genetic changes alone, we took a broader systems view to understand the underlying metabolic perturbations. We examined how the evolved variants rewired the fluxes through the major metabolic pathways that couple to the ETS. We generated RNA sequencing and metabolite profiling data for all the strains and performed targeted and systems-level analyses. We observed a high transcriptional correlation among the evolved replicates (Spearman’s rank correlation coefficient >0.75) of each variant, but the correlation between pre-and post-evolved variants was lower (Figure 2A). Notably, consistent genetic and transcriptomic changes supported a common evolutionary trajectory for the replicates of each variant.

**Figure 2:**
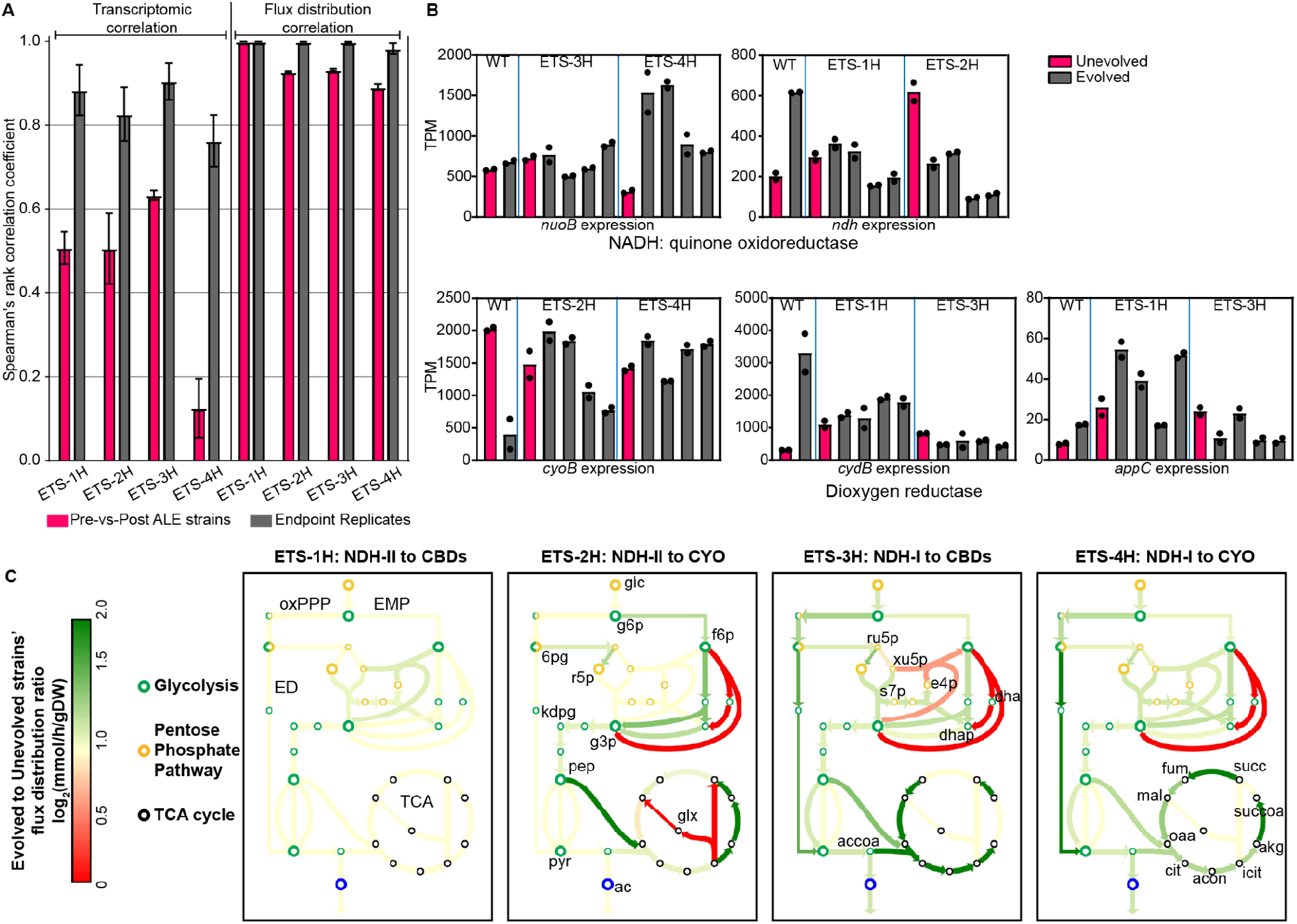
Metabolic rewiring supporting growth rate optimization in the ETS variants: (A) Transcriptional and metabolic flux distribution correlations between evolved replicates and unevolved strains. Error bars represent standard deviation. (B) Expression changes in the alternate NqRED and O_2_RED in the unbranched ETS variants. (C) Computed metabolic flux maps depicting the central metabolism in the evolved ETS variants as compared to respective unevolved ETS variants. Key metabolites are indicated in the figure as follows: glc, glucose; g6p, D-glucose-6-phosphate; f6p, D-fructose-6-phosphate; 6pg, 6-phospho-D-gluconate; ru5p, D-ribulose-5-phosphate; r5p, α-D-ribose 5-phosphate; xu5p, D-xylulose-5-phosphate; s7p, sedoheptulose-7-phosphate; e4p, D-erythrose-4-phosphate; dha, dihydroxyacetone; dhap, dihydroxyacetone-phosphate; kdpg, 2-keto-3-deoxy-6-phosphogluconate; g3p, glyceraldehyde-3-phosphate; pep, phosphoenolpyruvate; pyr, pyruvate; ac, acetate; accoa, acetyl-CoA; oaa, oxaloacetate; cit, citrate; acon, cis-aconitate; icit, isocitrate; akg, 2-oxoglutarate; succoa, succinyl-CoA; succ, succinate; fum, fumarate; mal, malate. [oxPPP, oxidative pentose phosphate pathway; EMP, Embden-Meyerhof-Parnas pathway; ED, Entner-Doudoroff pathway; TCA, Tricarboxylic acid cycle].

Bacterial physiology displays a remarkable compensatory potential facilitated by altered metabolic flux states resulting from genetic and transcriptomic changes ^22^. Therefore, we examined if the surrogate NqRED or O_2_RED compensated for the loss of function resulting from deleted ETS enzymes (Figure 2B). There was no clear compensatory trend in the strains with unbranched ETS except for ETS-4H. ETS-4H increased the expression of NDH-I while increasing or maintaining the expression of CYO after evolution. Surprisingly, the compensatory upregulation of *ndh* in *u*ETS-2H was lost after evolution to a higher growth rate.

Since RNA expression levels may not correlate with metabolic fluxes due to differential translation efficiency and different enzyme catalytic turnover rates, we performed a metabolic flux distribution analysis. To obtain the metabolic flux map, we measured the medium exchange rates of the major metabolites related to respiratory metabolism (Supplementary Table 2). We used both the metabolite exchange rates and transcriptomic data as constraints to simulate the flux through the pathways of the central carbon metabolism using a genome-scale model of metabolism and protein expression (ME-model) ^26^. We observed a high correlation in the metabolic flux distributions of the four evolved replicates of each strain, further supporting a similar evolutionary pathway followed by replicates of each variant (Figure 2A).

To more deeply understand the different metabolic states exhibited by the evolved variants, we examined the variations in their computed proteome allocation using the solutions from the phenotypic and transcriptomic constrained ME-models. We observed a clear distinction between strains with alternate NqRED for the preferred glycolytic pathway (Figure 2C). NDH-I has approximately 10-times higher molecular mass as compared to NDH-II ^27,28^. Therefore, despite its PMF generation potential, NDH-I is a less preferred dehydrogenase during oxic respiration to achieve faster growth ^5^. The non-proton pumping high turnover dehydrogenase, NDH-II, is better suited to relieve the growth bottleneck that may arise due to excess built-up of PMF while allowing the operation of oxic ETS ^5,29^.

The finite resource carrying capacity of a cell creates metabolic trade-offs on how to partition the proteome to support metabolic pathways best suited for a given growth condition. With an approximately 3.5-fold higher protein cost, the Embden–Meyerhoff–Parnass (EMP) pathway consumes a larger proportion of proteome as compared to the Entner–Doudoroff (ED) pathway ^30^. However, the higher ATP yield of the EMP pathway alludes to a potential tradeoff between the two glycolytic pathways for optimizing ATP production while maintaining a growth-supporting proteome ^31^. The ETS-3H and ETS-4H strains forced to respire using larger NqRED (NDH-I) increased the flux through the proteome conservative ED pathway. Thus, we observed a compensatory selection of the preferred pathway to achieve a balanced proteome.

Interestingly, while strains with *nuoB* deletion (ETS-1H and ETS-2H) increased metabolic flux through complex II of ETS, ETS-3H appeared to minimize the flux through complex II. Notably, *e*ETS-3H lacks *ndh* and acquired a mutation in the gene *sdhA* which codes for a complex II subunit. However, ETS-4H, which also lacks the *ndh* gene, increased the flux through complex II, albeit at a lower level compared to ETS-1H and ETS-2H.

Thus, metabolic plasticity (reflected in metabolic rewiring and associated proteome allocation) allows for redundancy in the *e*ETS variants while supporting the same growth rate. Knowledge of this metabolic plasticity motivated the examination of the overall bioenergetics-state of the evolved ETS variants to fully understand the basis for the evolution to the same growth rate. We have earlier defined an approach to classify the *E. coli* phenotypes into aero-types, which is a quantitative fitness descriptor based on cellular respiratory behavior and proteome allocation ^4^. The stratification of aero-types is based on the multimodal distribution of the fraction of total ATP produced through ATP synthase which is modulated through the discrete usage of ETS enzymes. We have reported a non-uniform distribution of phenotypic growth data in the rate-yield plane that can be approximately segregated in different aero-types based on sampling simulations. Here we used aero-types to examine the fitness distribution of ETS variants.

We observed that ETS-1H, ETS-2H, and ETS-3H did not show a major shift in their biomass yield during evolution and thus preserved their respective aero-types (Figure 3A). The evolutionary optimization of growth rate appears to be largely driven by rewiring central carbon metabolism while oxidative energy metabolism is conserved. ETS-4H jumped from a lower to a higher aero-type after evolution, suggesting an increase in oxic metabolism. The ETS-4H variant has the highest PMF generation capacity. Its aero-type shift to higher classes occurred only after adaptive evolution.

**Figure 3:**
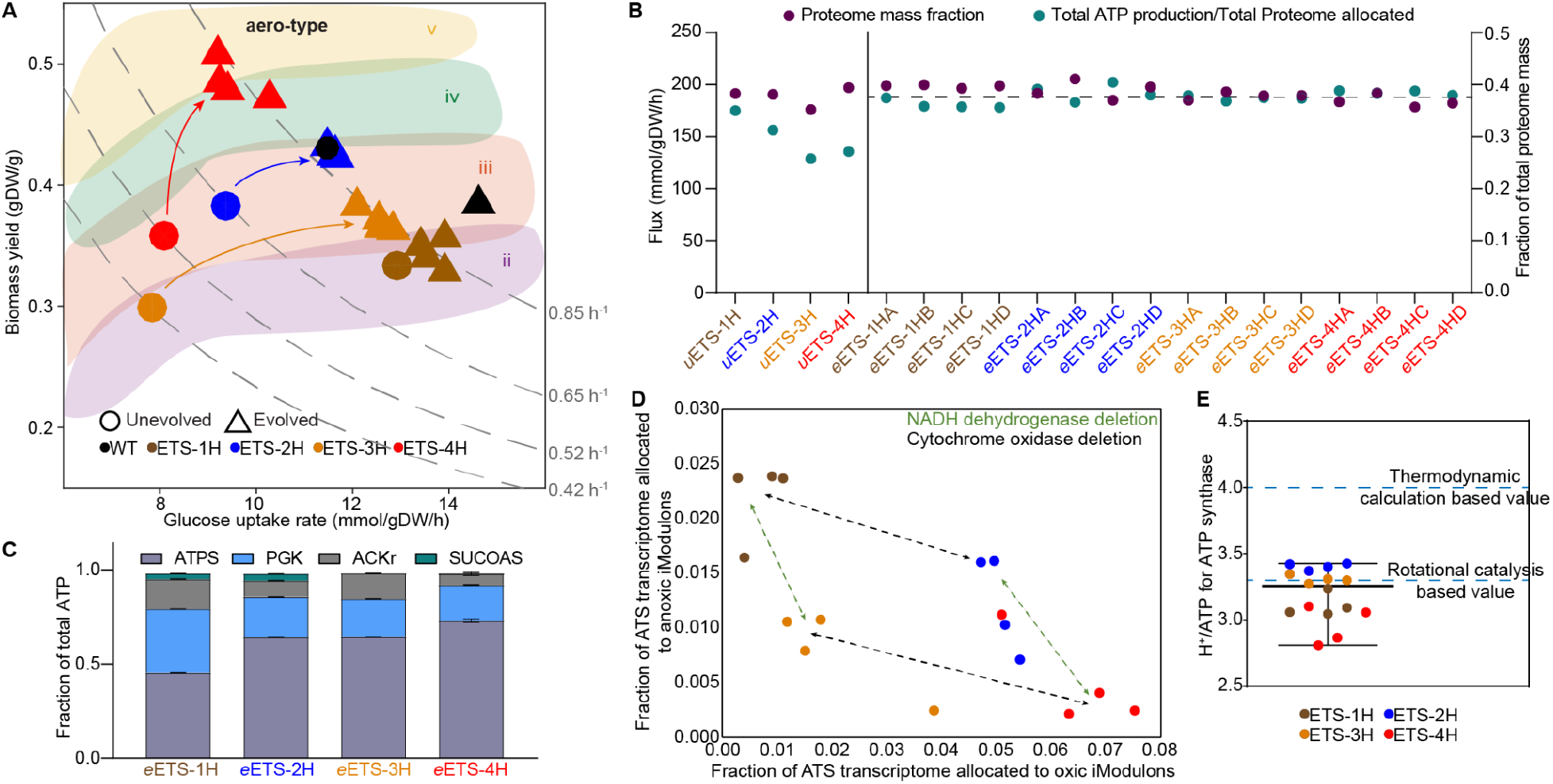
Systems-level examination of ETS variants: (A) Aero-type classification of the ETS variants. Broken lines on the aero-type plot show growth rate isoclines. (B) ME-model-based examination of the ATP production (left y-axis) and proteome allocation (right y-axis) in the ETS variants. The ATP produced per ATS proteome is approximately the same. (C) Contributions of different ATP-producing reactions towards total ATP production. ATP production by (i) ATP synthase (ATPS) in oxidative phosphorylation, (ii) acetate kinase (ACKr) in mixed acid fermentation, (iii) succinyl-CoA synthetase (SUCOAS) in the TCA cycle, and (iv) phosphoglycerate kinase (PGK) in the glycolysis pathway is shown in the histogram. (D) Tradeoffs in the expression levels of genes of iModulons associated with anoxic (y-axis) and oxic (x-axis) energetics underlie the rewiring of the ATS to allow all variants to achieve approximately the same growth rate. The lowest aerotype (ETS-1H) has high anoxic/low oxic gene expression while the highest aerotype (ETS-4) exhibits the opposite. The gene composition of the iModulons is shown in Supplementary Table 3. The outliers of the replicates for an ETS variant are reflections of the differences in their genotypes (Supplementary Figure 5). (E) Estimation of the number of protons required for the phosphorylation of ADP by ATP synthase (proton-to-ATP ratio) using the ME-model and the experimental data. The plot presents the median and range of values.

The clustering of each evolved ETS variant along the same growth rate isocline (Figure 3A) indicated global remodeling of the energy metabolic network to produce similar growth supporting bioenergetics. We thus defined a larger respiratory system, called the Aero-Type System (ATS), consisting of oxidative phosphorylation, glycolysis, pyruvate metabolism, the TCA cycle, and the Pentose Phosphate pathway, that together define the overall state of oxic energy metabolism (Supplementary Figure 2). The total proteome allocated to the ATS was very similar in each *e*ETS variant, and the total ATP output of each proteome expressed was almost constant (Figure 3B). Thus, the composition of the ATS was malleable and able to provide the same supply of ATP, allowing similar growth rates for all *e*ETS variants. We also observed a trend in the metabolic location of ATP production across the variants, where the relative contribution of oxidative phosphorylation was highest for *e*ETS-4H and lowest for *e*ETS-1H (Figure 3C). Accordingly, an inverse trend was observed for glycolytic and fermentative ATP production.

We next examined the transcriptome to identify the trade-offs in gene expression that enabled the different metabolic states. We applied a blind source signal separation algorithm, called independent component analysis (ICA)^32^, to examine differential partitioning of the transcriptome of the 209 ATS genes. ICA decomposed the ATS transcriptome into independently modulated sets of genes (called iModulons) (Supplementary Table 3). The activities of several iModulons showed a clear association with the aero-type of the ETS variants (Supplementary Figure 3). iModulons consisting of genes associated with oxic respiration showed a positive correlation with aero-type status (iModulons 8, 13, and b2287), and those constituted by anoxic and/or metabolic genes showed a negative correlation (iModulons 7, 9, 10, 16, and b3366) (Supplementary Figure 4). Thus, an oxic-anoxic transcriptomic trade-off enabled the four ETS variants to maintain similar ATP production capacity (Figure 3D).

With our comprehensive definition of the state of the ATS amongst the variants, we could address the issue of ATP synthase proton-to-ATP ratio. The number of protons translocated through ATP synthase to produce one molecule of ATP (H^+^/ATP) is still an area of active research. The rotational catalysis-based calculation suggests the H^+^/ATP value to be 3.3, due to the symmetry mismatch between the F_o_ and F_1_ complexes of ATP synthase: 3-fold symmetry of α3β3 in F1 and 10-fold symmetry of the c-ring in F_o_ ^33,34^. However, the proton-to-ATP ratio may vary depending upon any change in the number of c-subunits. The H^+^/ATP value derived using a synthetically reconstituted membrane system was found to be 4 ^35^. We used data generated on the variants to computationally estimate the most likely proton-to-ATP ratio for *E. coli* ATP synthase ^36^. We constrained the ME-model using the observed metabolic exchange rates and gene expression data and optimized for the H^+^/ATP value of ATP synthase that produces the experimentally estimated growth rates of the variants. The ME-model calculates the median value of the H^+^/ATP to be 3.25, a value close to 3.3 supporting the rotational catalysis hypothesis (Figure 3E). Notably, while 10 is the preferred number of *c* subunits in the F_o_ motor of ATP synthase, the number of subunits can vary, which will change the H^+^/ATP value ^37–39^.

Taken together, our results lead to an expanded definition of oxic respiration beyond the conventional ETS, which involves an electron transport chain to create PMF, that then drives the ATP synthase. Here, we define the Aero-Type System that encompasses the ETS and coupled metabolic pathways (Supplementary Figure 3). The ATS is composed of 209 genes. The ATS represents about 38% proteome allocation in all evolved variants. A decrease in the ETS energetic efficiency (often measured in terms of the P/O ratio) can be balanced by increased flux through the coupled metabolic pathways. This balance is governed by the cost of protein synthesis.

Remarkably, the overall proteome allocation to the ATS is similar in the evolved variants and generates the same amount of ATP, enabling them to achieve the same growth rate. The different ways in which the ATS is balanced underlies its plasticity and represents a demonstration of the key systems biology concept of alternate optimal states. These alternate states have a different combination of proton pumping efficiency, complementary metabolic rewiring achieved through tradeoffs in the composition of the transcriptome, and concomitant efficiency of proteome allocation, but enable the same overall cellular function. The cytoplasmic-periplasmic adaptive nexus that the ATS represents thus illustrates the deep plasticity inherent in achieving balanced energetic systems to match metabolic needs in different environmental niches.

## Acknowledgment

This work was funded by the Novo Nordisk Foundation Grant Numbers NNF10CC1016517 and NNF20CC0035580 and National Institutes of Health Grant R01GM057089. We would like to thank Marc Abrams (Systems Biology Research Group, University of California San Diego) for assistance with manuscript editing.

## Author contributions

A.A., A.F., and B.O.P. designed the study. A.A., C.O., and R.S. performed the experiments. A.A., A.P., K.C., P.P., C.L., and B.O.P. analyzed the data. A.A. and B.O.P. wrote the manuscript, with input from all co-authors.

