## Supplementary material for "Laboratory evolution of synthetic electron transport system variants reveals a larger metabolic respiratory system and its plasticity": Methods and Supplementary data

### Supplementary Information

**Contents:**

Supplementary Tables (3)

Supplementary Figure (4)

Methods

1) Examining PRECISE 2.0 for expression levels of respiratory enzymes
2) Strain generation and adaptive laboratory evolution
3) Prediction of the effect of amino acid substitutions
4) DNA resequencing and RNA sequencing
5) Phenotype characterization
6) Metabolic flux mapping and estimation of $H^+$/ATP value for ATP synthase
7) ATS proteome allocation calculation
8) ATS transcriptome ICA decomposition

Data Availability

Supplementary References

#### Supplementary Table 1: Key genetic changes in evolved ETS variants

| **Strain** | **Media adaptation-related mutations** | **Condition-specific convergent gene mutation** |
|---|---|---|
| *e*ETS-1HA | +5 bp between *hns* and *tdk* at genomic position 1293009, Δ82 bp between *pyrE* and *rph* at genomic position 3815859 | None |
| *e*ETS-1HB | +5 bp between *hns* and *tdk* at genomic position 1293009, Δ82 bp between *pyrE* and *rph* at genomic position 3815859 | None |
| *e*ETS-1HC | G to T substitution in *rpoC* at genomic position 4188513, Δ82 bp between *pyrE* and *rph* at genomic position 3815859 | None |
| *e*ETS-1HD | +5 bp between *hns* and *tdk* at genomic position 1292997, Δ82 bp between *pyrE* and *rph* at genomic position 3815859 | None |
| *e*ETS-2HA | A to T substitution in *rpoC* at genomic position 4187214 | None |
| *e*ETS-2HB | C to T substitution in *rpoC* at genomic position 4185540 | None |
| *e*ETS-2HC | A to G substitution in *rpoC* at genomic position 4187214 | None |
| *e*ETS-2HD | C to T substitution in *rpoC* at genomic position 4185540 | None |
| *e*ETS-3HA | Δ82 bp between *pyrE* and *rph* at genomic position 3815859 | T to C substitution in *sdhA* at genomic position 756099 |
| *e*ETS-3HB | Δ82 bp between *pyrE* and *rph* at genomic position 3815859 | C to T substitution in *sdhA* at genomic position 756150 |
| *e*ETS-3HC | Δ82 bp between *pyrE* and *rph* at genomic position 3815859 | T to G substitution in *sdhA* at genomic position 756886 |
| *e*ETS-3HD | Δ82 bp between *pyrE* and *rph* at genomic position 3815859 | C to G substitution in *sdhA* at genomic position 756968 |
| *e*ETS-4HA | C to A substitution in *rpoA* at genomic position 3440923 | C to T substitution in *yjjX* at genomic position 4633743 |
| *e*ETS-4HB | Δ3 bp in *rpoB* at genomic position 4183399 | C to T substitution in *yjjX* at genomic position 4633634 |
| *e*ETS-4HC | A to G substitution in *rpoC* at genomic position 4186578 | C to T substitution in *yjjX* at genomic position 4633657 |
| *e*ETS-4HD | C to T substitution in *rpoC* at genomic position 4189448 | C to T substitution in *yjjX* at genomic position 4633657 |

The intergenic mutation between *pyrE* and *rph* was present in the respective unevolved strain itself and these are listed here for a better context of media adaptation. Other mutations present in unevolved strain have not been listed here. The genes mutated in all four evolved replicates of a strain are listed here as convergent gene mutations.

**Supplementary Table 2:** Phenotypic characterization of the strains of the study

| Strain | Growth rate ($h^{-1}$) | Glucose uptake rate (mmol/gDCW/h) | Acetate secretion rate (mmol/gDCW/h) |
|---|---|---|---|
| *u*ETS-1H | 0.76, 0.79 | 12.77, 13.1 | 12.01, 12.35 |
| *e*ETS-1HA | 0.88, 0.9 | 13.57, 14.24 | 13.14, 13.61 |
| *e*ETS-1HB | 0.87, 0.82 | 13.61, 13.23 | 13.03, 12.88 |
| *e*ETS-1HC | 0.83, 0.82 | 13.84, 13.96 | 13.24, 12.93 |
| *e*ETS-1HD | 0.83, 0.83 | 13.35, 13.77 | 12.64, 13.06 |
| *u*ETS-2H | 0.65, 0.64 | 9.51, 9.26 | 6.41, 6.08 |
| *e*ETS-2HA | 0.87, 0.92 | 11.24, 11.71 | 7.76, 7.92 |
| *e*ETS-2HB | 0.9, 0.88 | 11.57, 11.67 | 7.77, 7.4 |
| *e*ETS-2HC | 0.9, 0.9 | 11.62, 11.68 | 7.36, 7.53 |
| *e*ETS-2HD | 0.89, 0.89 | 11.65, 11.76 | 7.98, 7.85 |
| *u*ETS-3H | 0.42, 0.43 | 7.68, 8.02 | 6.67, 6.51 |
| *e*ETS-3HA | 0.82, 0.83 | 12.6, 12.58 | 11.03, 11.26 |
| *e*ETS-3HB | 0.82, 0.86 | 12.35, 12.72 | 11.18, 11.26 |
| *e*ETS-3HC | 0.83, 0.84 | 12.78, 12.87 | 11.27, 10.97 |
| *e*ETS-3HD | 0.83, 0.84 | 11.92, 12.26 | 10.81, 11.07 |
| *u*ETS-4H | 0.53, 0.52 | 7.96, 8.18 | 3.95, 3.81 |
| *e*ETS-4HA | 0.85, 0.84 | 9.4, 9.01 | 4.24, 3.77 |
| *e*ETS-4HB | 0.86, 0.88 | 10.06, 10.47 | 5.25, 6.22 |
| *e*ETS-4HC | 0.79, 0.83 | 9.2, 9.3 | 6.14, 5.97 |
| *e*ETS-4HD | 0.8, 0.82 | 9.32, 9.49 | 4.96, 4.84 |

The values for two independent replicates are listed in the table.The levels of Succinate, Lactate, Formate, Ethanol, and Pyruvate were below the detection limit.

**Supplementary Table 3:** Description of the ATS iModulons

| S. No. | iModulon | Genes (negative coefficients in red) |
|---|---|---|
| 1 | iModulon-13 | *sdhB, cyoD, sdhD, cyoA, sdhA, sdhC, cyoC, fumA* |
| 2 | iModulon-b2287 | *nuoB* |
| 3 | iModulon-8 | *ndh, cyoB* |
| 4 | iModulon-11 | *cydB, appB, cyoB, appC, ndh* |
| 5 | iModulon-b3366 | *nirD* |
| 6 | iModulon-7 | *nirB, grcA* |
| 7 | iModulon-2 | *narI, narJ* |
| 8 | iModulon-1 | *yliI, cyoD* |
| 9 | iModulon-16 | *hyaB, hyaA, hyaC* |
| 10 | iModulon-9 | *hybO, hybA* |
| 11 | iModulon-10 | *tktB, adhP, fbaB, talA, poxB* |

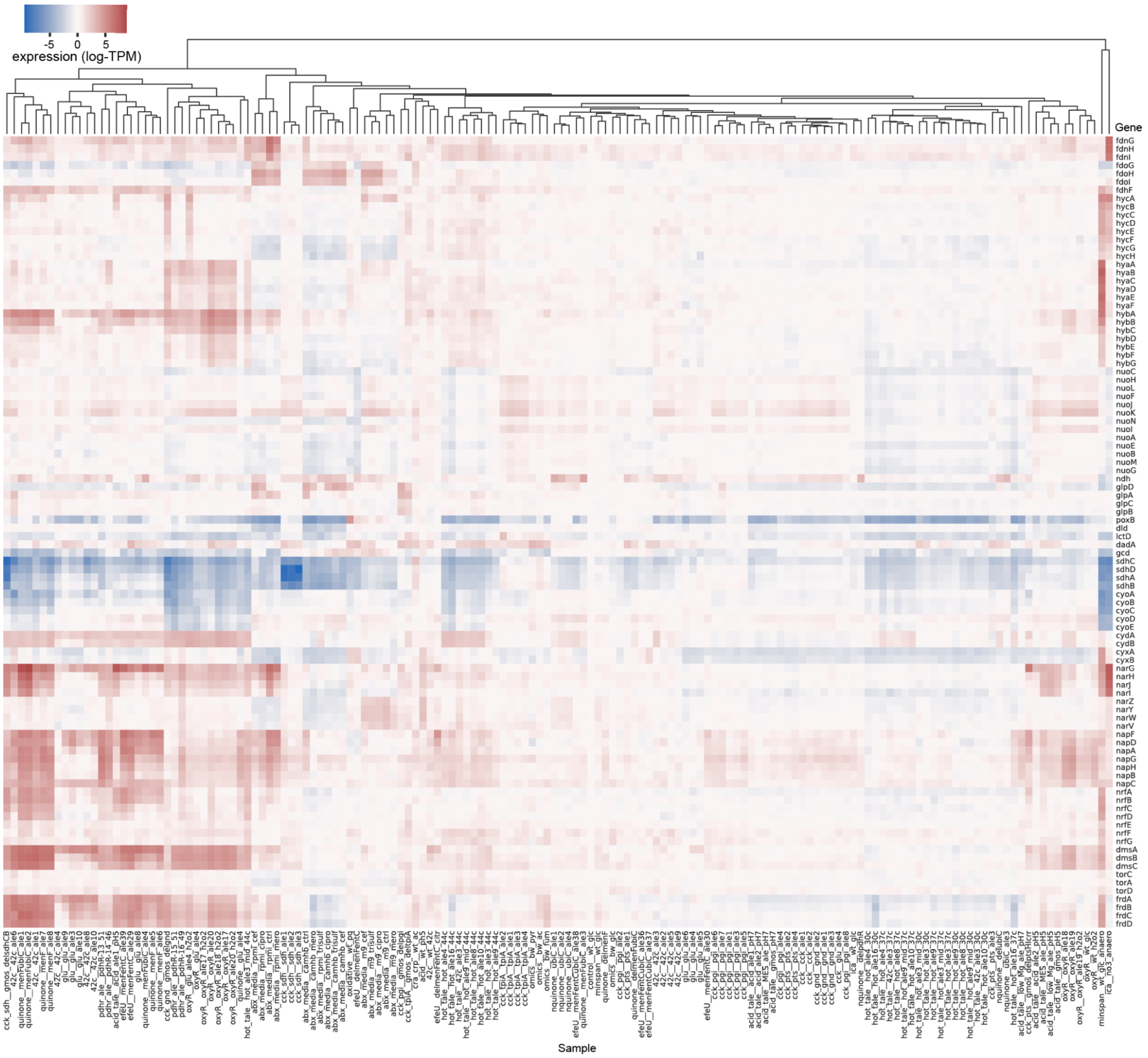


**Supplementary Figure 1: Expression levels of respiratory dehydrogenase and reductases of *E. coli*.** The entire RNA-seq compendium was explored and the conditions directly or indirectly associated with energy metabolism have been displayed.

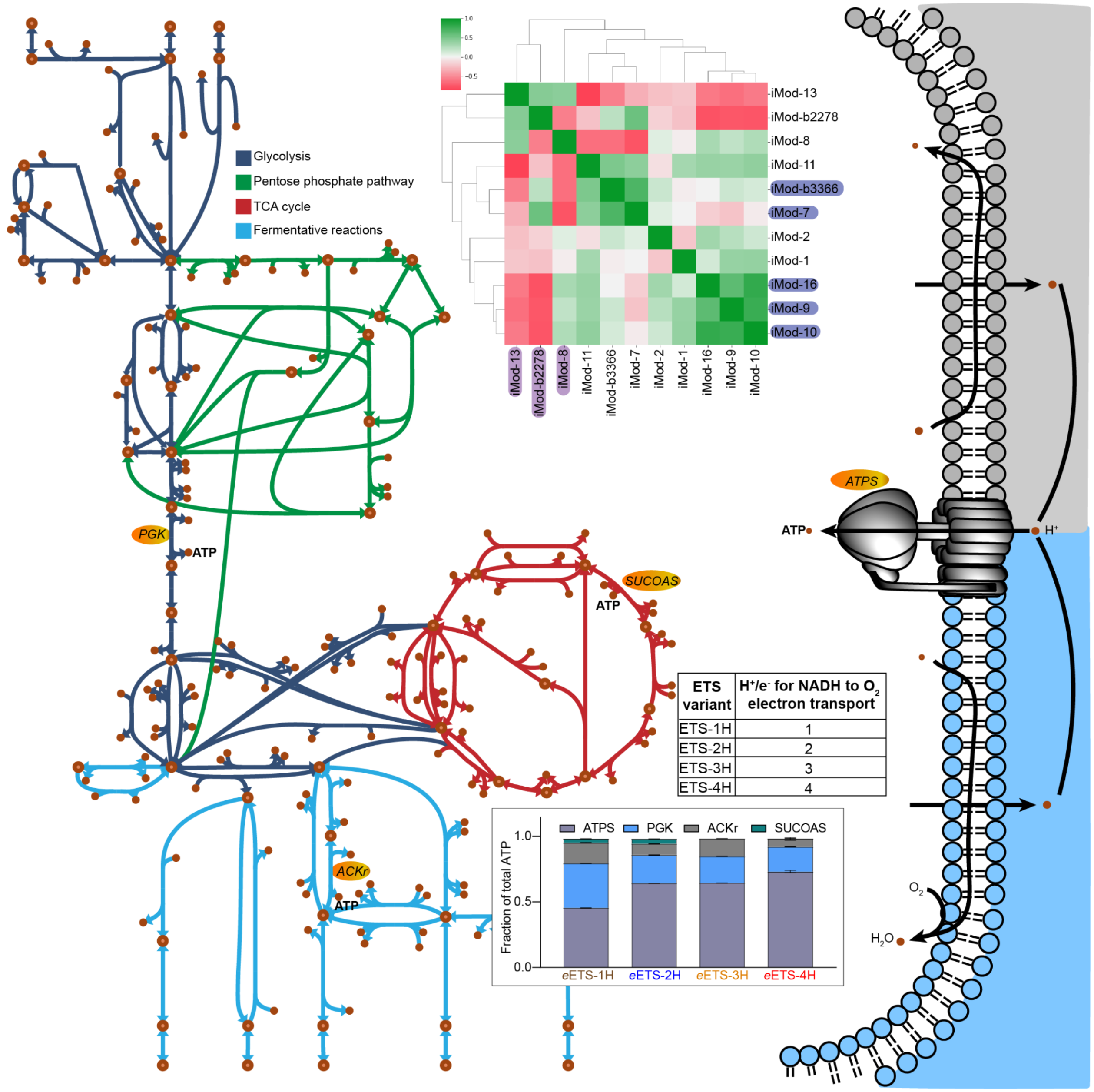


**Supplementary Figure 2: Scheme of Aero-type system (ATS)**. The pathways are showing reactions of ATS and major ATP production sites. Heatmap represents the correlation between the iModulons within ATS. iModulons showing a tradeoff are highlighted. The histogram from Figure 3 has been included to show adjustments within ATS to achieve a similar ATP yield. Part of the membrane highlighted in blue represents oxic ETS and another part highlighted in grey represents anoxic ETS. The list of ATS genes was generated based on COG and GO categories to include as many relevant genes as possible to represent pathways involved in ATP production, then filtered to remove genes that are never expressed in the multiple model simulations.

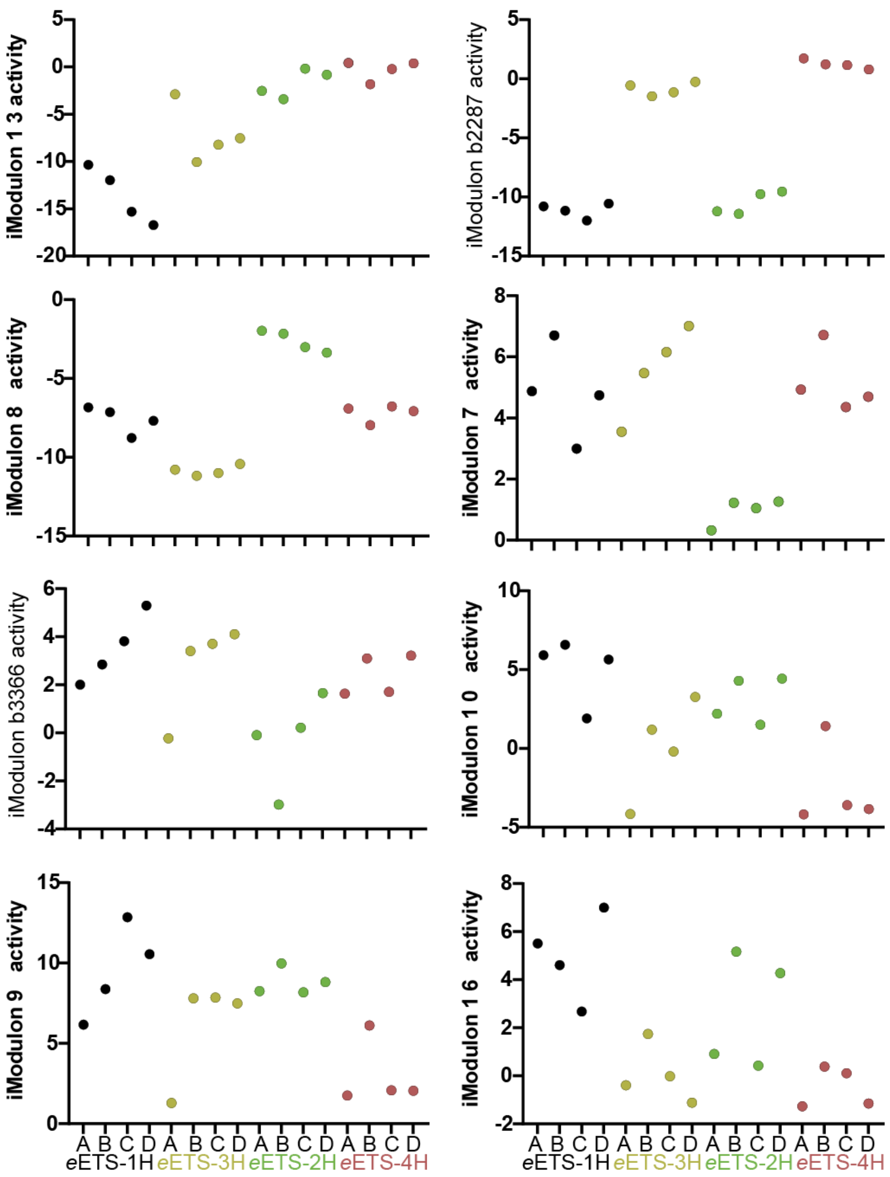


**Supplementary Figure 3:** Activities of ATS iModulons in the unevolved and evolved ETS variants calculated using independent component analysis. The gene membership of corresponding iModulons is listed in Supplementary Table 3.

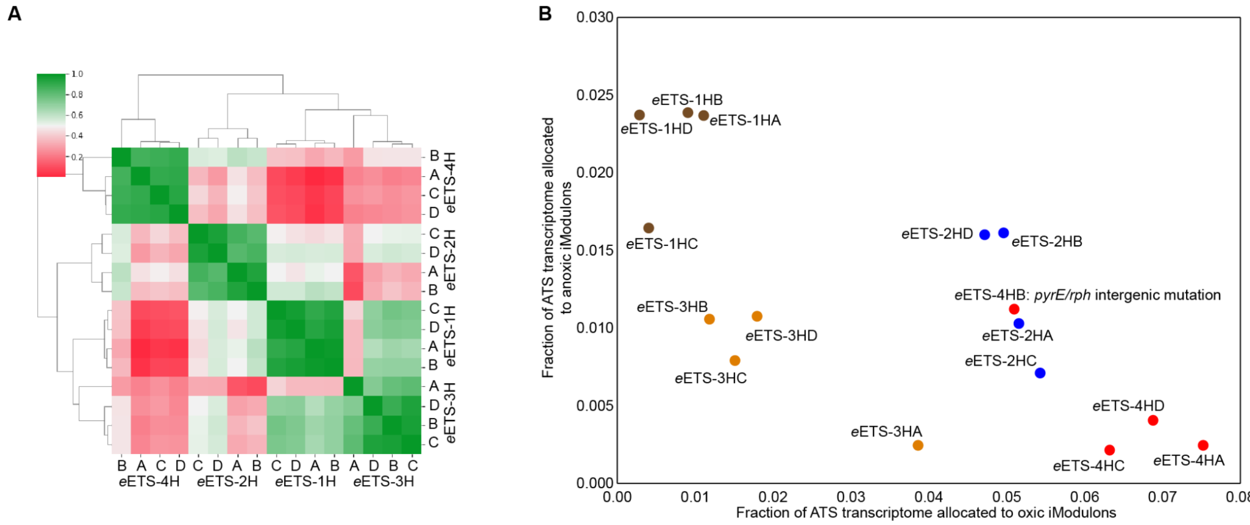


**Supplementary Figure 4:** (A) Correlation among ETS variants based on the ATS iModulon activities. (B) The enlarged representation of figure 3D lists the replicate information and potential genetic change in the outlier replicate *e*ETS-4HB.

### METHODS

#### 1.) Examining PRECISE 2.0 for expression levels of respiratory enzymes

PRECISE 2.0 is a compendium of high-quality RNA-seq for *E. coli* K-12 [1]. It contains 815 RNA-seq datasets of samples with different genetic changes or varied growth conditions. We examined the expression of respiratory dehydrogenases and reductases in the entire dataset. For intelligible purposes, we plotted the expression levels in samples that are directly or indirectly associated with energy metabolism. The expression levels shown are the median value across replicates for a sample.

#### 2.) Strain generation and adaptive laboratory evolution

*E. coli* K-12 MG1655 (ATCC 700926) was used as the wild-type strain. P1 phage transduction method was used to generate the knockout strains [2], and strains from Keio collection were used as a donor for the gene knockout cassettes[3]. *u*ETS-1H and *u*ETS-3H were generated and used for validation purposes in an earlier study [4]. *u*ETS-2H and *u*ETS-4H were generated here and all four ETS variants were evolved for this study.

ALE was performed using 4 independent replicates of each ETS variant. Cultures were serially propagated on M9 minimal medium with 4 g/L glucose at 37°C and well-mixed for proper aeration using an automated system that passed the cultures to fresh flasks once they had reached an $A_{600}$ of 0.3 (Tecan Sunrise plate reader, equivalent to an $A_{600}$ of ~1 on a traditional spectrophotometer with a 1 cm path length). Cultures were always maintained in excess nutrient conditions assessed by non-tapering exponential growth. The evolution was performed for a sufficient time interval to allow the cells to reach their fitness plateau.

**3.) Prediction of the effect of amino acid substitutions**

The ALE mutation datasets supporting the conclusions of this article is available in the following open-access archive repository: https://doi.org/10.5281/zenodo.5431595. These datasets are also available in the ALEdb database [5].

Mutated DNA sequence data processing was performed using Python 3. The mutations from ALEdb are described according to their experiment, evolution replicate, sample, and technical replicate. Some evolutions include midpoint samples that could inflate the frequency a mutation is observed. Unique ALE mutations were therefore only considered once per ALE. Starting strain mutations and hypermutator samples were filtered out of the ALE experiment mutation datasets according to their publications. Mutation needle plots were generated using the trackViewer R software package [6]. The visualizations for the 3D protein structures were generated using the *NGL* software package [7]. The software implementation of these actions is available in the following open-access archive repository: https://doi.org/10.5281/zenodo.5431595.

Mutation effects were predicted according to multiple methods. Truncations were predicted according to the potential effect of mutations on the function of start codons and their potential to introduce a premature stop codon. The predicted deleterious effects of SNPs were assumed according to significant SIFT (sorting intolerant from tolerant) scores (SIFT score < 0.05) [8]. The predicted structural destabilization effects of SNPs were assumed according to predicted significant ΔΔG scores (ΔΔG > 2)[9]. SIFT and ΔΔG scores were acquired from *Mutfunc*[10]. Functional annotations were acquired from *UniProt* [11] and *Mutfunc*. The software scripts supporting the prediction of mutation effects on the encoding of genes described in this article are available in the following open-access archive repository: https://doi.org/10.5281/zenodo.5431595.

**4.) DNA sequencing and RNA sequencing**

A clone from the endpoints of evolved strains was picked for DNA sequencing and RNA sequencing. The strains were grown in an M9 minimal medium supplemented with 4g/l glucose. Total DNA was sampled from an overnight grown culture and total RNA was sampled from a culture at an A600 ~0.6. Nucleic acid isolation, library preparation, and subsequent analysis were performed as previously described [12].

**5.) Phenotype characterization**

Phenotype characterization was performed using two independent biological replicates. Samples for the substrate uptake and secretion rate were collected at regular intervals and filtered using a 0.22 µm filter (PVDF, Millipore). The measurements were performed using refractive index detection by HPLC (Agilent 12600 Infinity) with a Bio-Rad Aminex HPX87-H ion exclusion column. The HPLC method was the following: injection volume of 10 µL and 5 mM H2SO4 mobile phase set to a flow rate and temperature of 0.5 mL/min and 45°C, respectively.

The phenotype dataset was used for the aero-type classification of the strains as described previously [4].

#### 6.) Metabolic flux mapping and estimation of $H^+$/ATP value for ATP synthase

Flux mapping were done as previously described using a genome-scale model of metabolism and protein expression [13]. The same FoldME model was used for estimating the $H^+$/ATP value for ATP synthase within each ETS variant and replicates. The model was constrained with phenotypic data (glucose uptake rate, acetate production rate) and expression data was layered on using the same methods used for the flux mapping [13]. In addition to these constraints, the necessary ETS genes for each variant were knocked out. Proton pumping ratios from 2.5 to 4.5 were sampled by changing the stoichiometry of the ATPS4rpp reaction in the ME-model, and then the proton pumping ratio was optimized so that the model produced a biomass dilution rate that matched the experimentally determined growth rate.

#### 7.) ATS proteome allocation calculation

The same FoldME model was used for the proteome allocation calculation as the flux mapping and ATP synthase estimation calculations. The model was constrained with phenotypic data (glucose uptake rate, acetate production rate, growth rate) and expression data was layered on using the same methods used for the flux mapping. Solutions from the fully constrained ME-models were then used for calculating proteome allocation. Total proteome allocation for each strain was calculated as follows:

$$Total\ Proteome\ Allocation\ =\ \sum_i mw_i * V_i^{translation}$$

Where $mw_i$ and $V_i^{translation}$ represents the molecular weight and translation flux of the *i*th protein in the model. Total proteome allocated to the ATS was calculated as follows:

$$Proteome\ Allocated\ to\ ATS\ =\ \sum_i mw_i * V_i^{translation}$$

where $mw_i$ and $V_i^{translation}$ represents the molecular weight and translation flux of the $i$th protein in the ATS (209 genes total). The list of 209 ATS genes was generated based on Clusters of Orthologous Groups (COG) and Gene Ontology (GO) categories to include as many relevant genes as possible to represent pathways involved in ATP production, then filtered to remove genes that are never expressed in the multiple model simulations. Mass fraction of proteome allocation to the ATS was calculated as a ratio of the two values for each strain.

Calculation of the total ATP produced by the ATS used the same fully constrained ME-model. A list of all metabolic reactions associated with ATS genes was curated. Reactions that consumed or produced ATP were noted and the stoichiometric coefficient associated with ATP was used as a modifier for calculating the total ATP production as follows:

$$Total\ ATP\ Production\ =\ \sum_i c_i * V_i^{metabolic}$$

where $c_i$ and $V_i^{metabolic}$ represents the ATP stoichiometric coefficient and the metabolic flux of the $i$th ATS associated reaction in the table below.

| **Reaction*** | PPK | PPK2 | ATPS4rpp | PFK_2 | HEX1 | PYK | PFK | PPS | PGK | GLGC | PFK_3 | GART | PPAKr | ACCOAL | ACS | ACKr | SUCOAS |
|---|---|---|---|---|---|---|---|---|---|---|---|---|---|---|---|---|---|
| **ATP coefficient** | -1 | -1 | 1 | -1 | -1 | 1 | -1 | -1 | -1 | -1 | -1 | -1 | 1 | -1 | -1 | -1 | -1 |

*Reaction IDs with corresponding reactions in the BiGG Database (bigg.ucsd.edu)

Total ATP Production/Total Proteome Allocated was calculated as a ratio of the total ATP production to the mass fraction of proteome allocated to the ATS for each strain.

#### 8.) ATS transcriptome ICA decomposition

Independent component analysis was performed on an RNA-seq dataset with steps described in [1]. The only genes included in the dataset were those contained in the list of 209 ATS genes. The dataset consisted of all unevolved strains, $u$ETS-1H through 4H, and all evolved replicates *e*ETS-1HA through *e*ETS-4HD. Additionally, the unevolved and evolved wild-type strains were included with the former being used as a reference to center the data. The final and resulting dataset that was used for ICA contained 209 genes by 22 conditions.

**DATA AVAILABILITY**

Resequencing and expression profiling data that support the findings of this study will be made available upon request. All ME-model-based simulations performed in this manuscript can be reproduced using the FoldME model, which is constructed using the COBRApy toolbox for constraint-based modeling and its extension for ME-models, COBRAme, ECOLIme, and solveME, all publicly available on Github (https://github.com/SBRG/cobrame/, https://github.com/SBRG/ecolime, https://github.com/SBRG/solvemepy). Biological materials of this work are part of our future work and we will make them available to the public after a year from the date of publication of this manuscript.
